# The structure and evolutionary diversity of the fungal E3-binding protein

**DOI:** 10.1101/2022.04.20.488913

**Authors:** Bjoern O. Forsberg

## Abstract

The pyruvate dehydrogenase complex (PDC) is a central metabolic enzyme in all living cells composed majorly of E1, E2, and E3. Tight coupling of their reactions makes each component essential, so that loss impacts oxidative metabolism pathologically. E3 retention is mediated by the E3-binding protein (E3BP), which has not previously been clearly resolved within the PDC. Here, the structure of the fungal E3BP in complex with the PDC core from *N*.*crassa* is resolved to 3.2Å, showing its mode of binding. Fungal and mammalian E3BP are shown to be orthologs, arguing E3BP as a broadly eukaryotic gene. Fungal E3BP architectures predicted from sequence data and computational models further bridge the evolutionary distance between *N. crassa* and humans, and suggest discriminants for E3-specificity. This is confirmed by similarities in their respective E3-binding domains, where a novel interaction is also predicted that may affect the interaction of its lipoyl substrate with recruited E3.

## Introduction

The 2-oxoacid dehydrogenase complexes are a class of protein complexes with acyl-transferase activity in metabolism and amino acid synthesis(Patel *et al*, 2014).They co-localize 3 catalytic components (E1-3) that act in sequence upon a substrate-carrying lipoyl domain(LD) that is flexibly linked to E2. This couples the reactions(Kato *et al*, 2008) and increases their overall rate through so-called substrate channeling. The pyruvate dehydrogenase complex (PDC) is the 2-oxoacid complex responsible for the bulk production of acetyl-coenzyme A (CoA) in all cells, crucial to oxidative phosphorylation, but also histone acetylation in the nucleus(Sutendra *et al*, 2014). Dysfunction and regulation of the PDC is consequently implicated in many disorders characterized by altered metabolism, including cancer(Sutendra & Michelakis, 2013; Guevara *et al*, 2017; Kaplon *et al*, 2013; Koukourakis *et al*, 2005). Genetic abnormalities also impact the PDC(Gray *et al*, 2014; Patel *et al*, 2012), and more recently it has also been recognized that the PDC is a source of reactive oxygen species, which impacts signaling cascades and the metabolic state of the cell(Fisher-wellman *et al*, 2013).

The arrangement of 2-oxoacid dehydrogenase complexes vary(Izard *et al*, 1999), but are invariably built around the C-terminal catalytic transacetylase domain (CTD) of E2. The CTD forms trimers that arrange into higher order assemblies. Further, E2 has two N-terminal domains separated by flexible linking regions. The N-terminal lipoyl domain (LD) shuttles the pyruvate-derived acetyl group via a covalent lipoamide modification. The central domain of E2 is a peripheral-subunit binding domain (PSBD) that tethers E1 to the CTD-composed core(Fig. 1A). It has been previously established that the mammalian PDC employs an E2 paralog to recruit E3 via a specialized PSBD named E3-binding protein (E3BP)(Powers-Greenwood *et al*, 1988; Rahmatullah *et al*, 1989; Marcucci & Lindsay, 1985). Mammalian E3BP has identical domain topology to E2, but is catalytically inactive (Harris *et al*, 1997), and has been concluded to substitute one or more E2 core units in an unknown fashion. Several models of the mammalian core E2:E3BP ratio have been suggested and studied(Brautigam *et al*, 2009; Hiromasa *et al*, 2004; Hezaveh *et al*, 2016, 2017, 2018; Maeng *et al*, 1996; Hackert *et al*, 1983; Vijayakrishnan *et al*, 2011), but no consensus has been reached, nor is the reason for the catalytic inactivity of E3BP known. Like mammals, fungi also utilize E3BP to recruit E3 (Maeng *et al*, 1994, 1996; Stoops *et al*, 1997). Unlike mammals however, fungal E3BP binds to the interior of the E2 core assembly instead of substituting core components(Forsberg *et al*, 2020; Skalidis *et al*, 2021; Tüting *et al*, 2021)(Fig. 1B). Mammalian and fungal E3BP have been treated as separate entities in literature, perhaps mainly due to the fundamental difference in their C-terminal domain and its mode of binding. The fungal E3BP has been denoted “protein X” (PX). Both E3BP and PX have influenced similarity-based automatic annotations, leading to confusion in current databases. To distinguish it from the CTD of E2 and reflect their shared features, the C-terminal domain of E3BP or PX is here named as the core-binding domain (CBD), regardless of taxonomic origin.

**Figure 1.**
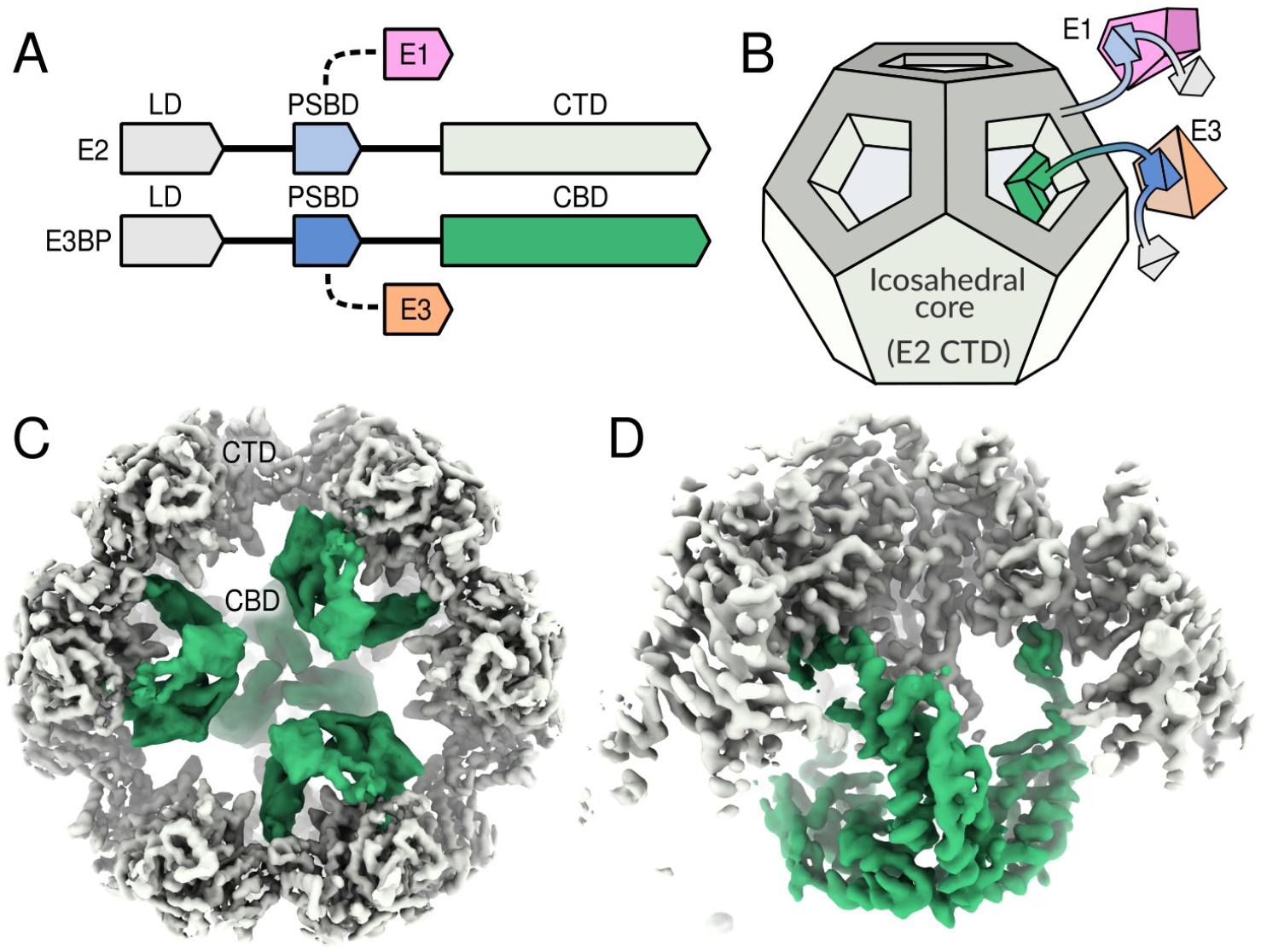
Cryo-EM reconstructions of *N. Crassa* E2:E3BP-CBD sub-complex. **A** Domain topology of E2 and E3BP, and **B** The overall arrangement of these protein in the fungal PDC. **C** The recombinant CTD:CBD sub-complex is reconstructed with tetrahedral symmetry to comply with a maximal CBD:CTD ratio, here colored as E3BP CBD (green) interior and E2 CTD core complex (gray). This permits symmetry expansion, signal subtraction, and further alignment and classification of CTD trimers. **D** The increased resolution of the core interior CTD trimer ultimately permits atomic modeling of the CBD trimer.

In this study, the molecular structure of the CTD:CBD subcomplex of the fungal PDC from *N*.*crassa* was determined using cryo-EM to 3.2Å resolution. It unambiguously demonstrates the determinants for their interaction, the mode of E3BP oligomerization interior to the E2 core, and the homology of the CBD and CTD. It is shown that neo-functionalization of E3BP from an ancestral E2 gene likely predates the reduction of the fungal CBD, which has subsequently diverged structurally from that of e.g. mammals, while preserving essential E3BP-functionality. Fungal and mammalian E3BP are thus determined as orthologs. This suggests that the E3BP function may be much more ubiquitous to eukaryotes than previously thought. In line with this, variations on the CBD are found in fungi outside the *Pezizomycota (Pez)* subphylum that *N*.*crassa* belongs to, among which the CBD appears either more or less similar to its ancestral CTD fold. In addition, the observed binding mode of fungal CBD is compatible with a much more complete E2-like fold, suggesting that core-internal localization is possible in non-fungal species. Sequence analysis of their respective PSBD corroborate the orthology of fungal and mammalian E3BP, and permits the first reliable analysis of its specificity for E3. In addition, the flexible linker connecting the *N*.*crassa* CBD and PSBD shows a conservation pattern that suggests relevance for interaction with the lipoyl domain (LD) as it interacts with E3. These findings comprehensively furthers our understanding of the separate recruitment of E3 to the PDC and its range of variation among eukaryotes, and suggests new modalities in this crucial metabolic complex.

## Results

### Overall structure

To determine the interactions of the fungal E3BP:E2 subcomplex, their CBD and CTD respectively were recombinantly co-expressed and examined by cryo-EM. Tetrahedral symmetry was used for the preliminary reconstruction(Fig. 1C), and 3-fold symmetric sub-complexes were computationally isolated by symmetry expansion and signal subtraction. CBD-occupied E2-trimers were selected by classification, and their distribution across PDC core particles analyzed. 4% of CBD trimers were assigned to E2 core particles with 5 or more CBD-trimers, which is not permissible from steric considerations(Forsberg *et al*, 2020). These are attributed to false positive identifaction, within the margin of error of data classification. 90% of PDC core particles had at least one CBD-trimer bound. Thus identified CBD-occupied E2 trimers were then further classified and aligned. This resulted in elevated CBD occupancy as evidenced by reconstructed intensity, and reduced conformational heterogeneity (flexibility). The final reconstruction to 3.2Å utilized less than one CBD-trimer per well-aligned PDC core particle (Fig. 1D), and permits the CBD from *N*.*crassa* E3BP to be built de novo.

The core fold of the *N*.*crassa* CBD consists of four helices and two flanking beta-strands (Fig. 2A), inherited from an ancestral CTD. The CBD of *N*.*crassa* has evolved to form a trimeric interface and symmetric axis that coincides with that of the core E2 CTD-trimer it binds to(Fig. 1D). This trimeric interface is unlike that of the CTD trimeric interface, and is instead evolved from the dimeric interface that core CTD trimers form to form larger assemblies. It is hydrophobic in nature, formed largely from the C-terminal end of the first CBD helix residues L285, I289, V291, P420, L423 and V424(Fig. 2D). R301 has clear side-chain density resolved (Fig. S1), and is highly conserved across *Pez* fungi. It likely stabilizes the C-terminal end of the same CBD monomer. The highly irregular C-terminal residues of the CBD in *N*.*crassa* is also noteworthy, since the penultimate residue is L/V/I in 95% of examined *Pez* sequences, whereas it is Arg only in *Neurospora*.

**Figure 2.**
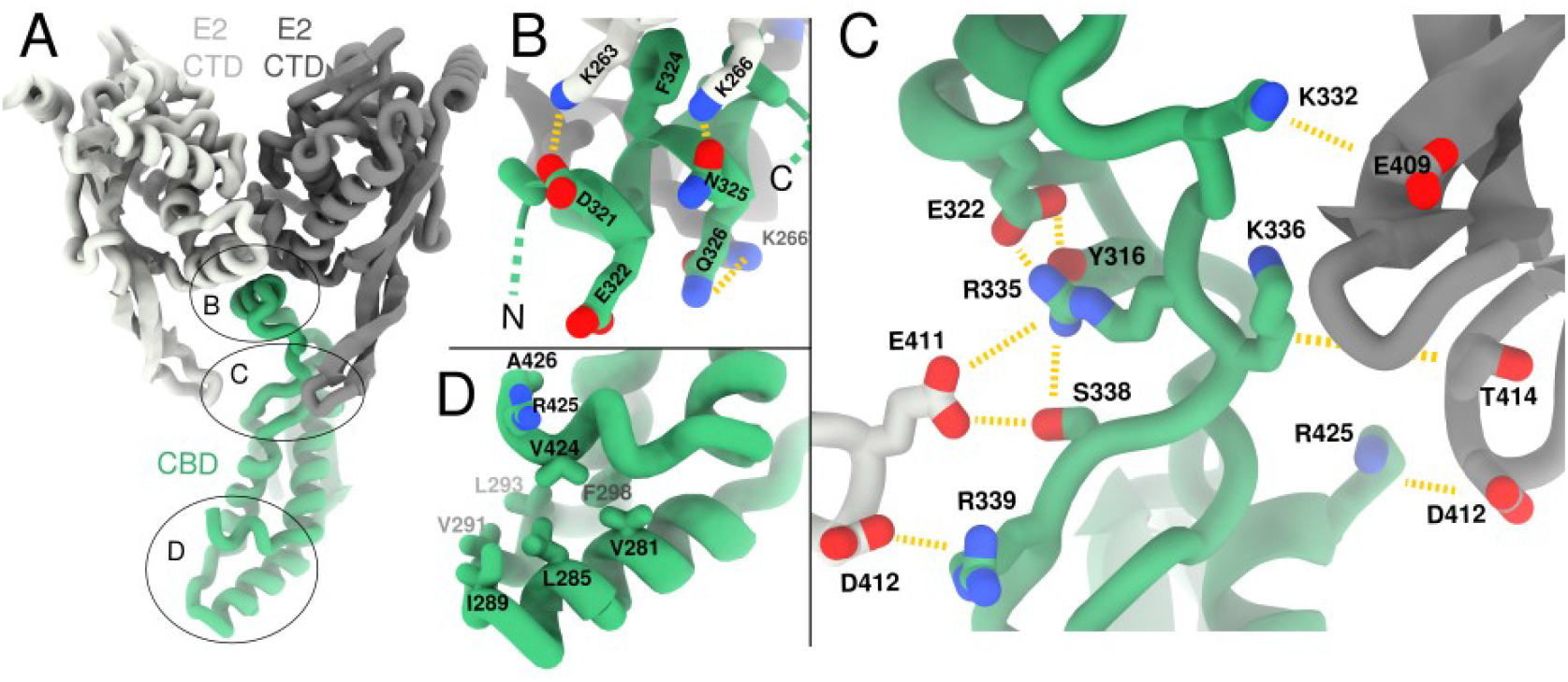
Molecular interactions and homology of fungal E3BP. **A** The E2 CTD forms trimers that further interact through a twofold-symmetric interface. Here, this dimer interface is shown, displaying one CTD from each of two such trimers in white and gray, respectively. The CBD of fungal E3BP binds in this interface, here showing a single CBD monomer. Indicated regions are shown in subsequent panels. **B** The binding motif is centered around a hydrophobic patch on the CTD complemented by CBD F324, and charge complementarity to CTD lysines K266 and K263. **C** Numerous potential salt bridges and electrostatic interactions are mediated by the E2 core-internal loop of up to three separate CTD monomers. **D** The CBD homomeric trimer interface is exclusively hydrophobic, barring R425, which is unresolved and atypical of the fungal CBD.

Beyond the core fold, the CBD contains a coil region (S338-S348) that lines the helices adjacent to the beta-sheet where Y342 shields the hydrophobic core of the CBD and may interact with E401 and R404. One would expect F346 and possibly L385 to pack against I299 and F393, but this hydrophobic patch appears incompletely shielded from solvent. Hence, this appears to be a partially stabilized, flexible loop. This flexible loop precedes one of two extended regions that protrude from the CBD core fold, each containing a small and highly conserved motif denoted M2 (T319-L328) and M3 (D360-A367) respectively (Forsberg *et al*, 2020). M2 is a clearly resolved short helix connected to the CBD by structured loops, and is responsible for binding to E2 (Fig. 2B). M3 is flanked by regions of predicted disorder 347-289, none of which is resolved in the current reconstruction. The termini of the M3 disordered region are within a 5Å of each other, and positioned near the 5-fold pseudo-symmetry axis of the E2 core assembly, placing it approximately equidistant from each of the 5 closest E2 trimers (Fig. S2A).

### E2-E3BP interactions

As shown previously(Forsberg *et al*, 2020; Tüting *et al*, 2021), the CTD-trimers form a homomeric dimer interface and hydrophobic pocket to which the CBD of fungal E3BP binds (Fig. 2B). The present reconstruction shows that M2, and not M3, is the major binding interface of the E2 core assembly. In *N*.*crassa*, M2 is composed of a amphipathic helix which is conserved within *Pez*, and which likely extends to *Saccharomycetes (Sac)* (Fig. 3B). A conserved Phe (*N*.*crassa* F494) appears crucial by protruding directly into the binding site, as predicted in *C. thermophilum(Tüting et al, 2021)*. Additionally, K266 from both participating CTD monomers interact with M2 residues N325 and Q326 from the CBD (Fig. 2B). K263 of the central E2 trimer is also poised to interact with CBD residue D321. As CBD binding is oriented wihtin the strictly symmetric pocket, binding of monomeric CBD is likely unstable. CBD trimerization provides avidity of binding in this scenario, and enforces oriented M2 binding. The M3 motif is not resolved in the present reconstruction, since it is part of the disordered region of *N*.*crassa* CBD connecting F346 and E391 (45aa). M3 however recapitulates the general properties of the de facto binding motif M2 (Fig. 3C). One must consider that M3 might act as a secondary binding motif against the CTD binding pocket. Residual density is observed in these interfaces (Fig. S2B), but cannot be clearly resolved. This density could alternatively be meta-stably bound M2 of monomeric CBD that does not form CBD trimers. The universal conservation of M3 thus remains unexplained.

**Figure 3.**
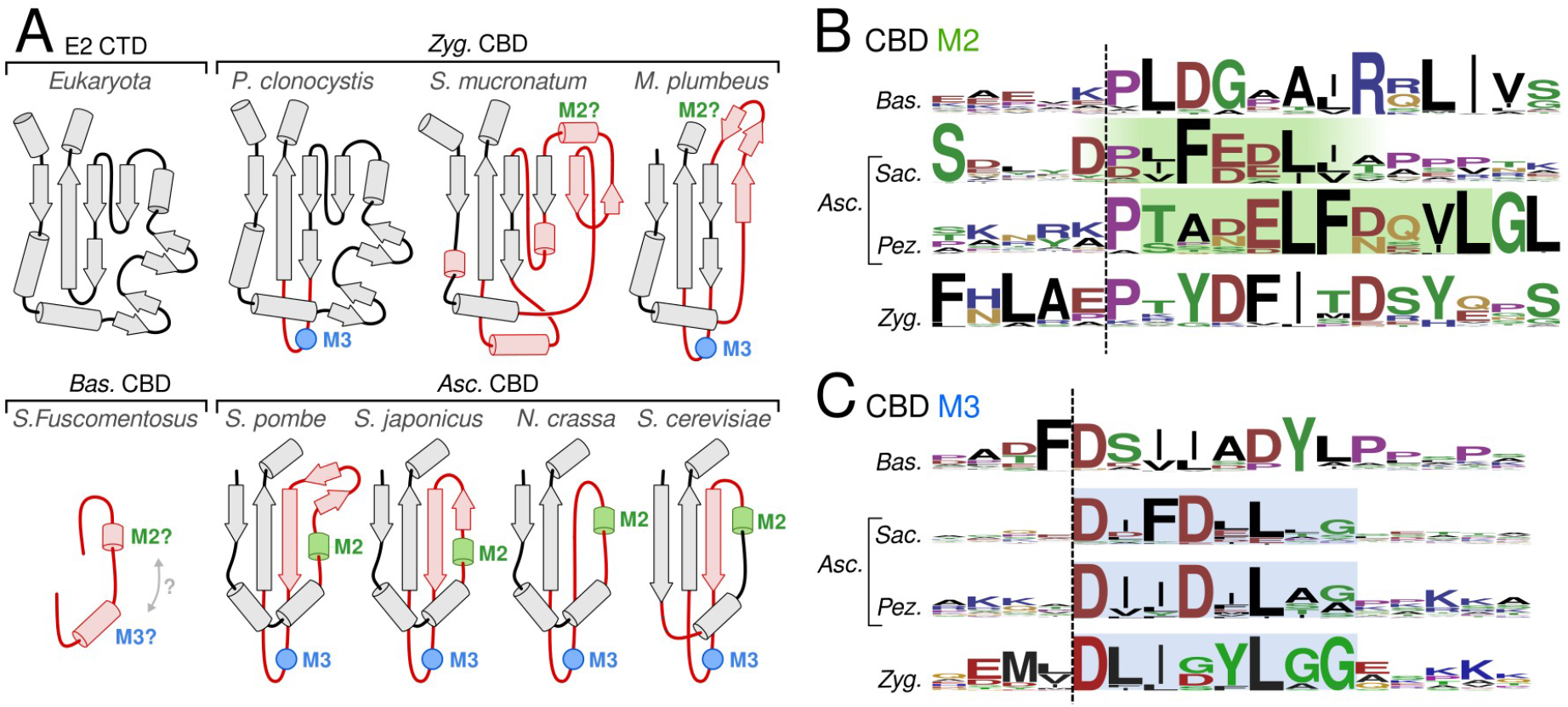
Topology of fungal E3BP. **A** The E2 CTD fold topology schematic of eukaryotic E2 CTD, compared to that of the CBD of E3BP in several fungal species, grouped by phylum. Topologies are sequence-based predictions apart from that of *N*.*crassa*, and display a broad variation in evolution from the ancestral CTD of a duplicated E2 gene. Substantial differences to the CTD is indicated in red, and conservation motifs are also indicated as M2 and M3 in *Ascomycota* (*Asc*). Inferred M2 and M3 sequence motifs in *Zygomyceta* (*Zyg*) and *Basidiomycota* (*Bas*) are indicated, with varying confidence. **B-C** The sequence logo of M2 and M3 regions are given for each fungal CBD phylum, and confident assignment is highlighted as based on inferred sequence similarity and conservation compared to *N*.*crassa*.

Flexibility in the CBD is evident from data processing(Fig. S3). There are however stabilizing interactions from the core-internal beta-strand loop of the CTD (Fig. 2C). R339_CBD is likely to interact with D412_CTD of the central trimer, and R335_CBD may also interact with E411_CTD. R335_CBD is also poised to stabilize the loop connecting M2 with the CBD core fold, through E322_CBD, Y316_CBD, and possibly also by proxy of S338_CBD (Fig. 2C). Additional CTD monomers may also form interaction pairs with the CBD through its core-internal loop: K332_CBD-E409_CTD, K336_CBD-T414_CTD and K403_CBD-D412_CTD (Fig. 2C). The CTD beta-loop supplying these potential interactions is universal to 2-oxoacid dehydrogenase complexes, but varies in length and composition across species and E2 substrate-specificity. No significance of the core-internal E2 beta-loop has been suggested, but here shows electrostatic stabilization to a core-internal binding partner.

### Animal and fungal E3BP are orthologs

E3BP and E2 are homologous based on similarities in domain topology. Computational modeling of *C. Thermophilum* E3BP also predicted that its CBD was a partial E2 fold(Tüting *et al*, 2021). The reconstruction of the *N*.*crassa* CBD validates this model and confirms that fungal E3BP arose by neo-functionalization of a duplicated 2-oxoacid acetyltransferase gene in fungi as well as in mammals. Orthology between mammal and fungal E3BP is not established, and E3BP has instead been described as a mammalian gene in literature(Jiang *et al*, 2018). The occurrence of E3BP fungi has not been demonstrated outside *Ascomycota (Asc)*, whereas orthology would imply a much broader occurrence. To affirm that fungal and mammalian E3BP are divergent orthologs as opposed to convergent through evolution, the characteristics of E3BP were generalized and used to query other fungi and eukaryotes through existing sequence databases. E3BP was identified as proteins displaying i) a domain topology consistent with E2 homology, ii) an E3-specific PSBD (IPR004167, IPR036625, CDD397105, SSF47005), iii) a catalytically inactive acetyltransferase-homologous CBD, and/or iv) an M3-like motif of unknown function. Three distinct forms of fungal E3BP were thus identified in existing databases. First, the E3BP of *Basidiomycota* (*Bas*) is inferred from criteria i) and iv). The CBD of *Bas* is reduced to two short stretches of predicted helical structure, which resemble M2 and/or M3 motif (Fig. 3B-C). In line with this, the inferred *Bas* CTD binding pocket is similar to that of *Asc*, with conservation of residues equivalent to *N*.*crassa* K263 and K266. This suggests that avidity through oligomerization as in *Asc*, without a steric limit to E3BP binding. Alternatively, dual binding motifs per CBD monomer may be utilized to increase binding affinity and occupy two binding pockets per CBD monomer. The *Bas* CBD also holds a short conserved ‘Y/F-L/F-DGLϕ’-motif (ϕ=hydrophobic). Second, the CBD of *Asc* mathces all characteristics i)-iv), since it includes those studied here and previously in *S. cerevisiae(Stoops et al, 1997), N*.*crassa(Forsberg et al, 2020)*, and *C. thermophilum(Tüting et al, 2021)*. Its CBD is a CTD-derived fold that is broadly similar to that of *N*.*crassa*(Fig. 3A). There are however variations in *Taphrinomycotina* and *Saccharomycotina* that will require structural confirmation to verify e.g. CBD oligomeric state. M2 and M3 can however be unambigously assigned thorughout *Asc*, based on similarity to *N*.*crassa*. Third, the CBD of *Zygomyceta* (*Zyg*) is also match all criteria i)-iv), but is far less consistent than *Asc*. Sequence data for *Zyg* is also more sparse than for *Asc*, but frequently display a CBD that is more similar in size and structural composition to its CTD ancestor(Fig. 3A). Some species display a locus and motif similar to M3, but that of M2 is less certain. Combined with the more CTD-like fold, it is unclear if the *Zyg* CBD will form core-interior binding or core-substitution. Were core-substitution possible, its core-internal loop is much longer than that of any mammalian counterpart (typically ∼50aa). It is also this loop that presents M3-like sequence features(Fig. 3C). Hence, *Zyg* E3BP clearly shows distinctively fungal properties, but is quite dissimilar from any *Asc* species. The tripartite division (*Bas/Asc/Zyg*) of fungi outlines the extent of fungal variation of the CBD as detected by the established search criteria(Fig. S6). The range of fungal CBDs and the shared features validate their shared origin and thus recapitulate the evolutionary history of the reduced CBD in *Asc*. It is therefore posed that fungal and animal E3BP are likely orthologs, with a shared evolutionary origin. This would argue E3BP is a eukaryotic gene. Indeed, numerous annotations as E3BP exist in eukaryotes outside mammals inferred from sequence analysis, but none that display the reduced fold apparent in fungi.

### PSBD specificity

To further validate the orthology of fungal and mammalian E3BP, and to characterize E1/E3 specificity, E2 and E3BP protein sequences of fungal phyla and animals were analyzed and compared within the PSBD. Multiple-sequence alignments (MSAs) of each (sub)phylum was established, and anlyzed in the context of several existing structures. All such structures agree that a PSBD is universally composed of two helices connected by a loop that collectively bury a hydrophobic core. Crystallographic structures of human E3BP-E3 complex (Ciszak *et al*, 2006; Brautigam *et al*, 2006, 2011), and the E2-E1 complex from a bacterial species(Frank *et al*, 2005; Pei *et al*, 2008) also suggest a singular binding interface and pose of the PSBD, regardless of taxonomy or specificity for E1 or E3. This is corroborated by further studies of bacterial PDC, where the PSBD of E2 is not selective towards E1 or E3(Chandrasekhar *et al*, 2013). To further aid sequence analysis, alphafold2(Evans *et al*, 2021) was utilized to establish multimer-folded structural models of human and *N*.*crassa* PSBDs from E2 and E3BP, interacting with E1 or E3. These recapitulate the expected binding pose of E2-E1 and E3BP-E3 complexes confidently (as judged by pLDDT score). Conversely, E2-E3 and E3BP-E1 complexes could not be confidently modeled, as expected. The model of the human E3-E3BP interface does not depart significantly from either 1ZY8(Ciszak *et al*, 2006), or 2F5Z(Brautigam *et al*, 2006). Previous sequence analysis posed that I157 of human E3BP provides specificity for E3(Ciszak *et al*, 2006), contrasted against R383 in the human E2 (Fig. 4A-B, position 38). The present analysis confirms this position as distinguishing of E2 from that of E3BP in animals, but the limited sequence variation within animals makes its significance questionable. In further evidence against I157 as directly mediateing E3 specificity, fungal species deviate from this pattern and cross-phylum analysis instead reveals that a single arginine on either side of the immediately preceding glycine is universally preserved. This pattern extends to animal sequences, where E3BP has a conserved arginine (Fig. 4A, position 38, human:R155). Hence, I157 alone does not supply specificity. Instead, three more consistently distinguishing sequence patterns that are common to both animal and fungal species are found, which have not been previously noted. The first such sequence pattern resides in the C-terminal end of the first helix of the E2 PSBD. (Fig. 4A, position 19-21). This polar motif has consensus sequence ‘EKG’, and is least prominent in *Zyg*. Similar residues are frequently found in E3BP, but conservation is only found in E2. The second notable sequence pattern is instead conserved in E3BP but not E2, and consists a hydrophobic motif in the C-terminal end of the second helix(Fig. 4B, position 45-48), following a universally conserved aspartate (Fig. 4B, position 43, human:D215, *N*.*crassa*:D206). Here, hydrophobic residues and leucine in particular are clearly preferred in E3BP. As part of this motif, a universal leucine (Fig. 4B, position 45, human:L217, *N*.*crassa*:L208) is exposed to solvent without any apparent shielding, as is a conserved hydrophobic reside 3 residues downstream (Fig. 4B, position 48, human:V220, *N*.*crassa*:L211). Like the polar E2 motif, it unambiguously discriminates the E3BP PSBD from that of E2 across both fungi and animals. The third pattern is lead by a universally conserved glycine that terminates the second helix in the E3BP PSBD. This termination motif has ‘GxI’ as consensus sequence (Fig. 4B, position 49-51, 54-56 in animals), where the isoleucine appears to stabilize the hydrophobic collapse of the PSBD fold. Whereas human I228 folds back onto the PSBD itself(Fig. 5B), *N*.*crassa* I214 fold back towards the previously mentioned and highly conserved L208. The termination motif has not been resolved structurally, as 2F5Z(Brautigam *et al*, 2006) includes residues up to E230 but can only resolve residues until T225. No available complexed structure includes the termination motif, whereas 2F60(Brautigam *et al*, 2006) finds residues up to E230 to form a continuous helix in the absence of E3 binding. All three aformentioned motifs discrimnate E2 from E3BP across both fungi and animals, and are thus in clear support of a shared origin and E3BP orthology across eukaryota. Curiously however, none of these motifs contribute to interactions in the canonical binding interface(Fig. S5A). Despite the clear significance of these motifs, present models and evidence cannot substantiate a rationale for their involvement in E3 specificity. One may additionally observe a fourth motif (Fig. 4B, position 51-67) that is spefici to *Asc*, separated from the canonical PSBD by a stretch of low conservation. To examine if this motif offers a further rationale to E3 specificity, the *N*.*crassa* E3-PSBD complex was computationally predicted wit hthis motif included. In this model, the region of low conservation from N215 to S230 (Fig. 4B, position 51-67) forms a third and amphipathic helix that lines one E3 monomer, extending towards its substrate pocket (Fig. 5B). The conserved motif following it is consistently and confidently modeled as a coil of alternating hydrophobic residues that line the E3 substrate pocket(Fig. 5B). The same structure, interactions, or confidence cannot be attributed to a human model, where the linking region is notably shorter and has a much higher proline content(Fig. 4B). These additional interactions are likely specific to the E3BP-E3 interaction and will contribute to specificity. It may thus be speculated that interactions beyond the canoncial interface may similarly contribute to E3 specificity through the three aformentioned conservation motifs that discriminate E2 from E3BP across eukaryota.

**Figure 4:**
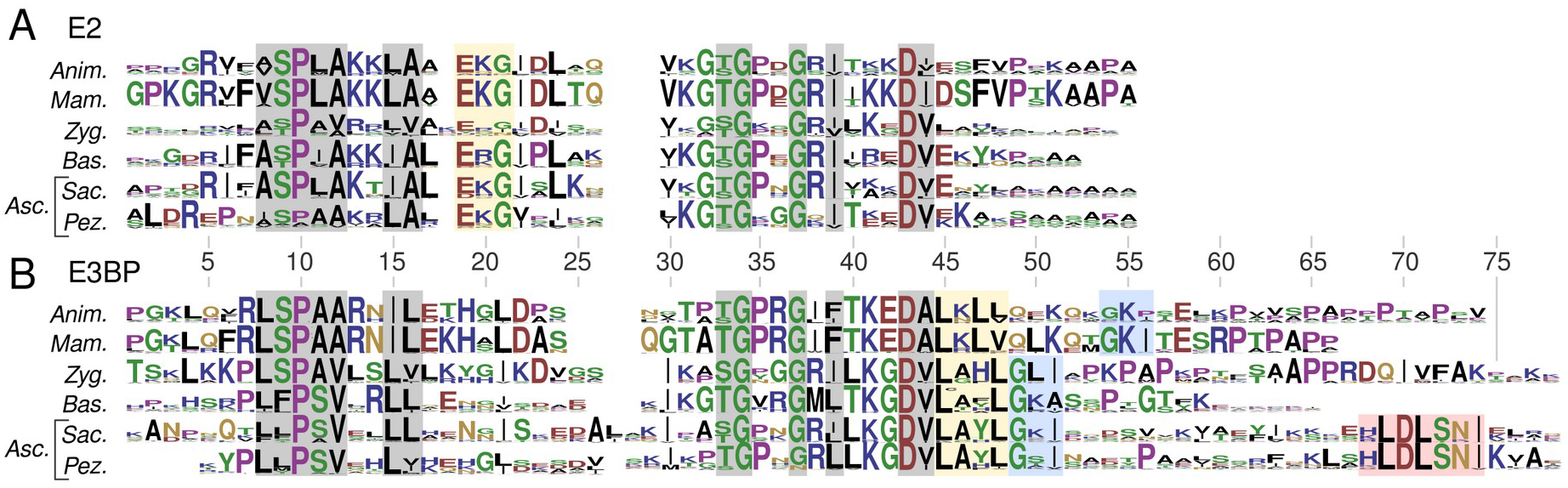
PSBD sequence logos of E2 and E3BP. **A** PSBD sequence alignment pertaining to E2p (pyruvate dehydrogenase transacetylase) from animals, mammals, and fungal phyla. **B** Alignment as in A, pertaining to E3BP. Regions are highlighted as universal to any PSBD (gray), discriminating E2p from E3BP in all examined groups (yellow), PSBD helix 2 termination motif in E3BP (blue), and predicted as extended PSBD E3-binding motif (red). The latter is lead by residue H231 in *N*.*crassa* (cf. Fig. 5).

**Figure 5:**
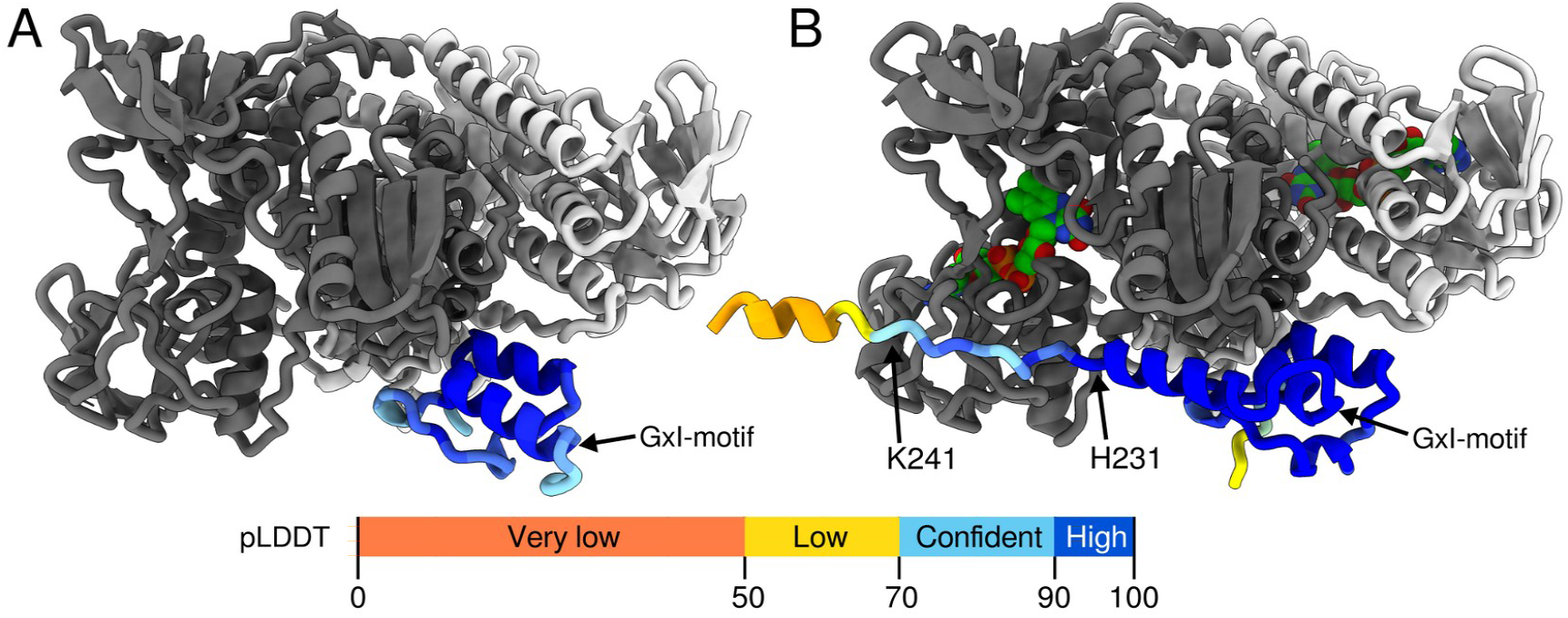
PSBD binding to E3. **A** Computational prediction of human E3BP:E3 complex (supp. data AF2.HS.E3-E3BP). Only the PSBD is colored by pLDDT, since all E3 residues shown have pLDDT>90. E3 monomers are individually colored by white and gray for clarity. The FAD cofactor molecule is shown (green). This model agrees well with present structures and models of binding. **B** Computational prediction of *N. crassa* E3BP:E3 complex (supp. data AF2.NC.E3-E3BP), colored as in A. Downstream of the GxI motif a helical region extends towards the E3 substrate pocket, which is further lined by E3BP residues H231-K241.

## Discussion

The cryo-EM reconstruction of the CBD from *N*.*crassa* E3BP inside the PDC core here reveals its oligomeric form and specific interactions. This permits a confident analysis of variations in the CBD across fungi, which in turn substantiates its orthology to human E3BP. This is further corroborated by sequence motifs in the PSDB that dicriminate E2 from E3BP in both fungi and animals. Given that E3BP in fungi and mammals is thus concluded to have descended from the same ancestral gene of their last common ancestor, all species subsequently diverged from this common ancestor could be expected to utilize an E3BP as part of its PDC. E3BP should thus be considered a eukaryotic gene, and annotations as “pyruvate dehydrogenase protein X component” should reasonably be universally changed to E3BP, since share a genetic origin. Moreover, they are proven to share E3-binding as primary function, and utilize their CBD to bind the E2 CTD, meriting this annotation.

Given it’s early occurrence in metabolic evolution, it is pertinent to ask why ancestral E3BP neo-functionalization from an E2 gene was favorable. Naively, recruitment of both E1 and E3 by a shared PSBD as in bacteria implies that their binding ratio is entirely determined by their relative affinity and availability. Utilization of E2 and E3BP as presently discussed decouples the recruitment of peripheral components E1 and E3. This offers a further rationale of why the fungi evolved to add rather than substitute PDC components. Through CBD addition (as opposed to substitution), the fungal PDC maintains a fixed capacity to recruit E1 through the E2 PSBD. Moreover, the 30 binding sites for E3BP impose a fixed E3 capacity as well, which in *N*.*crassa* is reduced to 12 through steric restraints imposed by the CBD trimer. The fungal PDC thus combines both decoupled and fixed stoichiometry of E1 and E3 recruitment. It stands to reason that less drastic changes to a core-substituting CBD might similarly dictate the assembly stoichiometry through heteromeric interfaces or decreased stability, as is suggested in the human PDC. This however remains an unproven hypothesis, whereas the fungal PDC is proven to utilize mechanisms to confine the component ratios.

What then drove CBD fold reduction as seen in present-day fungi? First, one must conclude that significant reduction of CBD must lead to its inability to substitute E2 CTD core subunits. Consequently, CBD fold reduction in fungi is only likely to have occurred after a mode of core-addition, such as M2, had been established. Once it had, one may speculate that fold reduction was necessary to permit entry into the E2 core interior. As a corollary question, could the full ancestral CBD oligomerize interior to the PDC? Curiously, the full E2 CTD can in fact be superposed on the fungal CBD trimer pose without major clashes. Remarkably, such a core-interior trimer would not even impose any further steric restraint to E3BP binding stoichiometry compared to that of e.g. *N*.*crassa* - 12 CBD monomers could still be accommodated inside the icosahedral E2 core(Fig. S6). In such an arrangement, beta-loop of the core-internalized CBD monomers would also extend towards each other and offer domain-swapping interactions(Fig. S6). Of note, the same loop in the CBD evolved to hold the unresolved M3 motif in the presently resolved CBD. It would thus be highly interesting to observe the core-interior arrangement of a CBD which is more akin to its CTD ancestor, such as e.g. *S. mucronatum* or *C. reveresa*. Importantly, the possibility of accommodating a full CTD in the alternate trimerization mode (as in *N*.*crassa* CBD) within the PDC core suggests that while CBD fold reduction likely favors core-internalization, it may not be necessary. The variations in CBD size and components implied by available fungal sequences could then simply correlate with germline mutation rate or environmental factors, but may also reflect the positive enforcement of binding avidity and binding stoichiometry provided through steric occlusion of the PDC interior. There is thus a clear motivation for the both the CBD addition model and CBD fold reduction, while neither is strictly necessary. The addition model also rationalizes the catalytic inactivation of the fungal CBD, whereas it is far less clear why the CBD is catalytically inactive in the proposed substitution model of mammals. This interesting parallel remains to be clarified by future research.

A question that similarly remains unanswered, regards the function of the M3 motif. It is contained within a region of predicted disorder, and is not resolved by the present reconstruction. It is however remarkably conserved throughout fungi. Is M3 an alternate binding motif? The oriented binding of M2 in the symmetric pocket indicates reversible binding that then presumably requires additional binding affinity. In the *N. crassa*, the CBD oligomer provides avidity, but one can also imagine that M3 provides additive affinity. *Pez* M3 is however more similar to *Zyg* M3 that to its own M2, which indicates that M3 is conserved by binding to an interface that is different to that of M2. Moreover, previous findings clearly indicate that M3 alone is insufficient to enrich E2 cores by affinity purification(Forsberg *et al*, 2020). Nevertheless, residual density is observed in the regions of E2 where E3BP would bind were it not sterically prohibited(Fig. S2), which could then be due to M3. Can the M3 loop reach these interfaces? The 44-residue loop begins and terminates 45-50Å away from the 3 closest E2-dimer interfaces that are not occupied by E3BP trimer M2. A fully extended protein can reach as far as 3.4Å per residue. WIthin the M3-containing loop, only 12 residues precede it and 25 residues follow it. The M3 motif therefore has a reach of only 40-45Å, unless the CBD partially unfold to lend it further reach. It is thus possible but unlikely that M3 binds to unoccupied dimeric CTD interfaces. The present reconstruction however offers some evidence to the contrary, as one would expect a maximally extended such structure to be less variable and thus resolved. There is no evidence of this. Additionally, such a narrow reach would result in oriented binding and one might similarly expect relatively well-resolved density, which is not observed. Taken together, this indicates that M3 binds weakly or non-specifically to the CTD pocket in the present reconstruction, whereas its physiological target is something else. Is there any evidence for M3 function in sequence data? In other species of fungi, such as *Sac*, the equivalent of M3 frequently contains a conserved phenylalanine, distinguishing it from *Pez* and making it more similar to *Pez* M2 (Fig. 3B-C). In S*ac*, the CBD also lacks fundamental structural elements of *Pez*, and thus it has not been directly observed to form trimers. Hence, the aformentioned additive affinity of M2 and M3 might be more reasonable in *Sac* than oligomer-mediate avidity as in *Pez*. Additionally, the trimeric form of *N*.*crassa* CBD provides a steric limit to 12 bound copies of E3BP per core assembly. In the absence of such steric restraints, additive affinty may similarly enforce a limit of 15 bound copies of E3BP in *Sac*. Alternatively, M3 might be responsible for binding E3BP to something other than PDC E2, such as another E2 dehydrogenase complex. It may also interact with some core-internal substrate or cofactor such as coenzyme-A. This is substantiated by the observation that the CBD of fungal species of *Zyg* may well be core-substituting as proposed for mammals, but still contain an extraordinarily long core-internal loop with a motif reminiscent of M3 in the same structural element. Clearly, the purpose of M3 is an interesting but outstanding question.

Finally, the present analysis suggests notable similarities in the PSBD of fungal and animal E3BP, and their mutual differences to that of E2. This solidifies the orthology of fungal and animal E3BP, and suggests discriminants for E3-binding. The polar motif of E2 and hydrophobic motif of E3BP are both situated on the same face of a solvent-exposed region of the PSBD, albeit on different helices. Because existing structures and established models do not suggest any direct interactions with these residues, an altered PSBD binding pose would be required to argue their direct responsibility for binding specificity for either E1 or E3. This is certainly possible, however the available sequence data shows overwhelming support for the established mode of binding. Instead, one might consider that this interface offers an opportunity for secondary or even tertiary binding. In such an event, the functionality may also be something other than E1/E3 specificity. An example of this is provided by the novel structural element presently predicted in the E3-PSBD complex of *N*.*crassa* (Fig. 5B). This feature is unique to *Asc*, and does not implicate the E3 specificity through the polar or hydrophobic motifs within the PSBD, but serves as an example of a possibly important interaction interface that has so far eluded our understanding. Future research will be necessary to elaborate on the polar, hydrophobic, and termination motifs within thePSBD of both E2 and E3BP. In further reference to the predicted structural element and interaction of the extended *Asc* PSBD (*N*.*crassa* H231-K241), one might speculate that it affects the binding the lipoyl domain to E3. If so, it would potentially differentiate the two available binding pockets available on each E3 dimer, as only one PSBD binds each such dimer. The implications of this observation are interesting, but will need experimental confirmation. Regardless, the universal conservation of this motif within *Asc* exemplifies that the flexible linkers of 2-oxoacid dehydrogenases are not as passive as previously assumed. In the absence of more specific alternate hypotheses, present evidence indicates that it augments binding and/or specificity for E3 in *Asc*. Similar conservation is not found outside *Asc*. Its conservation is highly interesting in itself, as it constitutes substantial differences between the fungal and human E3BP, while one still finds its PSBD and CBD to contain highly conserved and defining features that cannot currently be assigned a function. Further investigation will be necessary co clarify these interesting correlatives and discrepancies.

## Materials and methods

### Sample preparation

Sample was prepared as previously described(Forsberg *et al*, 2020). Briefly, bacterial co-expression of *N. crassa* E2 (residues 225–458, uniprot:P20285) and his-tagged E3BP (residues 261–426, uniprot: Q7RWS2) utilized the petDuet dual-expression vector. The vector was amplified using *Escherichia coli* DH5-α, and expressed in Rosetta2 (DE3). For expression, cells were grown in terrific broth at 37C and 180 r.p.m. until OD reached ∼0.5, then induced by 1mM final IPTG. Cells were harvested 3 h post induction. Cells were pelleted and re-suspended in 50 mM Tris pH7.5, 0.5 M NaCl, 20mM imidazole, then lysed by high-pressure homogenization. Intact cells and debris were pelleted, and the supernatant collected. Ni-NTA agarose slurry was added and incubation under agitation proceeded for at least 30 min. Isolation and washing of Ni-NTA was performed by gravity flow column, and eluted with 300mM imidazole. Imidazole was exchanged by size-exclusion chromatography (SEC) on a GE Superose 6-increase 3.2/100, and spin-concentrated

### Data collection

Grids for cryo-EM were prepared by glow discharge in a Pelco easiGlow. 3 μl of sample was applied to 300-mesh 1.2/1.3 quantifoil grid and vitrified in a FEI Vitrobot mark IV, following 30 s wait, 2 s blot, and 2 s additional wait before plunging. 100% humidity and 4C was maintained prior to plunging. 4063 Micrographs were collected on an 300kV FEI Krios with a Gatan K2-GIF in counting mode, at a nominal maknification of 165k (0.86Å/px). At total dose of 31.4 e/Å^2^ was fractionated across 32 frames over 4s. Grid screening and optimization, as well as data collection was conducted at the Swedish National Cryo-EM Facility at SciLifeLab, Stockholm University and Umeå University

### Data processing

In order to resolve fungal E3BP better than previously possible, single-particle cryo-EM data was reprocessed in RELION(Zivanov *et al*, 2018) using a procedure essentially identical to that of localized reconstruction(Ilca *et al*, 2015) by symmetry expansion and classification. In brief, PDC core particles were aligned and reconstructed under icosahedral symmetry. CTF-refinement and polishing were iterated to refine aberration parameters. PDC core particles were next reconstructed by tetrahedral symmetry to aid subsequent C3-reconstruction. The tetrahedral reconstruction was the basis for symmetry-expansion around each icoshaderal 3-fold symmetry axis, to align each E2 trimer of every icosahedral particle to the same reconstructed E2-trimer. As a result, subtraction of complementary density and re-sizing images at the equivalent position of this E2 trimer permitted additional alignment and classification of E2-trimers pertaining to every original icosahedral particle under C3 symmetry. More details are provided in supporting Fig. S3.

### Model building

The cryo-EM half-maps of the limited PDC sub-complex were used to sharpen the full map using DeepEMhancer(Sanchez-garcia *et al*),and residues 265-346 and 390-425 of *N*.*crassa* E3BP were build *de novo* against the using coot(Casañal *et al*, 2020). The coordinates of E2 previously published (PDB:6ZLO) were used as a starting point for the modeling of E2. Two E2 monomers forming the dimeric binding pocket were included to permit the full extent of E2-E3BP interactions to be modeled. The complex was threefold expanded to model homomeric E3BP interactions, and refined in Phenix(Liebschner *et al*, 2019) as a threefold symmetric complex. As the focused cryo-EM reconstruction following subtraction and re-centering did not encompass the E2 monomer adjacent to the central E2 monomer, some of the models was not supported by density during refinement. Negligible asymmetric effects were observed due to this, but B-factors are consequently higher than expected. The full adjacent monomer was nevertheless retained in the deposited structure. Figures were visualized using ChimeraX(Goddard *et al*, 2018) and PyMOL(Schrödinger & DeLano, 2020).

### Bioinformatics and sequence alignment

BLAST(Boratyn *et al*, 2013) was utilized to search for E3BP homologs as per the established criteria. Jalview(Waterhouse *et al*, 2009) was used to manage sequence data, aligned using the clustal(Madeira *et al*, 2019) alignment algorithm. Sequence sets were frequently pruned manually to omit sequences which did not inform on the sought features. It should therefore be noted that possibly erroneous annotations or extremely divergent sequence data may not be included. No filtering for sequence redundancy was performed, but transcripts annotated as ‘partial’ were removed. For animal sequences, E3BP was discriminated from E2 based on catalytic inactivity as inferred by the absence of a histidine in its canonical transferase triad. For fungal species, sequences were discriminated as E3BP based on mutual similarity in the CBD and the established criteria (see text and methods).

## Acknowledgments

The cryo-EM data were collected at the Swedish national cryo-EM facility, staffed by M. Carroni, J. M. de la Rosa Trevin, J. Conrad, and S. Fleischmann. The manuscript drew benefit from critical evaluation by Pranav Shah. The work was funded by the Swedish research council.

## Data availability

The atomic model of the E2-E3BP complex was deposited in the PDB (7R5M) and the map of the limited cryo-EM reconstruction deposited in the EMDB(14331). The EMDB entry provides the full map, as well as the half-maps, mask, and map sharpened by DeepEMhancer(Sanchez-garcia *et al*). All established MSAs can be accessed <u>here</u>, and all computational models are available <u>here</u>.

## Author contributions

B.F. Collected and analysed data, and wrote the article.

