## Supporting Tables and Figures for "The structure and evolutionary diversity of the fungal E3-binding protein"

### Supplementary table 1

Reconstruction and model parameters

| | | | |
|---|---|---|---|
| **EMDB identifier** | 14331 | | |
| **Symmetry** | C3 | | |
| **Sharpening factor** | -133 | masked | unmasked |
| **Resolution [Å]** | half-map FSC 0.143 | 3.3 | 3.6 |
| | sharpened FSC 0.143 | 3.2 | - |
| | d99 (f/h1/h2) | 4.3/1.8/1.8 | 4.2/1.8/1.7 |
| | map-model FSC (f/h1/h2) | 2.6/3.0/3.4 | 3.0/3.2/3.8 |
| **PDB identifier** | 7R5M | | |
| **Composition** | Chains | 9 (6x E2 + 3x E3BP) | |
| | Atoms | 27096 | |
| | Hydrogens | 13761 | |
| | Protein: | 1758 | |
| | Water | 0 | |
| | Ligands | 0 | |
| **RMSD** | length [Å] (#>4sig) | 0.003 (0) | |
| | angles [°] (#>4sig) | 0.613 (0) | |
| **Isotropic B-factors** | min | 30 | |
| | mean | 84.17 | |
| | max | 206.99 | |
| **Molprobity** | | 1.37 | |
| **Clashscore** | | 3.1 | |
| **Ramachandran** | Outliers | 0% | |
| | Allowed | 3.23% | |
| | Favored | 96.77% | |
| **Rotamer outliers** | | 1.22% | |

### Supplementary table 2

List of computationally modelled proteins

| Accession | Protein(s) | Annot | Species | Subkingdom | Division/phylum | Subdivision/subphylum | Class |
|---|---|---|---|---|---|---|---|
| RMX77418 | E3BP | hypo | Hortaea werneckii | Dikarya | Ascomycota | Pezizomycotina | Dothideomycetes |
| XP_017996629 | PX | hypo | Phialophora attinorum | Dikarya | Ascomycota | Pezizomycotina | Eurotiomycetes |
| XP_018001728 | PX | PX | Phialophora attinorum | Dikarya | Ascomycota | Pezizomycotina | Eurotiomycetes |
| XP044666154 | PX / PDXK | unchar | Bacidia gigantensis | Dikarya | Ascomycota | Pezizomycotina | Lecanoromycetes |
| PBP15521 | E2? | PX | Diplocarpon rosae | Dikarya | Ascomycota | Pezizomycotina | Leotiomycetes |
| KAF3097484 | PX | pyridox | Orbilia oligospora | Dikarya | Ascomycota | Pezizomycotina | Orbiliomycetes |
| RPA78695 | PX | hypo | Ascobolus immersus | Dikarya | Ascomycota | Pezizomycotina | Pezizomycetes |
| RPB21680 | PX | PX | Terfezia boudieri | Dikarya | Ascomycota | Pezizomycotina | Pezizomycetes |
| XP_956161 | PX | PX | Neurospora crassa | Dikarya | Ascomycota | Pezizomycotina | Sordariomycetes |
| Q5AKV6 | PX | hypo | Candida albicans | Dikarya | Ascomycota | Saccharomycotina | Saccharomycetes |
| P16451 | PX | PX | Saccharomyces cerevisiae | Dikarya | Ascomycota | Saccharomycotina | Saccharomycetes |
| XP019021831 | PX | unchar | Saitoella Complicata | Dikarya | Ascomycota | Taphrinomycotina | incertae sedis |
| XP018226045 | PX | hypo | Pneumocystis carinii | Dikarya | Ascomycota | Taphrinomycotina | Pneumocystidomycetes |
| O94709 | PX | PX | Schizosaccharomyces pombe | Dikarya | Ascomycota | Taphrinomycotina | Schizosaccharomycetes |
| XP_002172135 | PX | PX | Schizosaccharomyces japonicus | Dikarya | Ascomycota | Taphrinomycotina | Schizosaccharomycetes |
| XP041229627 | PX | PX | Suillus Fuscotomentosus | Dikarya | Basisomycota | Agaricomycotina | Agaricomycetes |
| XP022780480 | PX | PX | Stylophora pistillata | non-fungi | Cnidaria | | Hexacorallia |
| KAG9290904 | PX | hypo | Geosiphon pyriformis | Zygomyceta | Glomeromycota | Glomeromycotina | Glomerales |
| CAG8600220 | PX | - | Paraglomus brasilianum | Zygomyceta | Glomeromycota | Glomeromycotina | Paraglomerales |
| XP0055108998 | ? | PX | Aplysia californica | non-fungi | Mollusca | | Gastropoda |
| KAG0028626 | PX | hypo | Podila clonocystis | Zygomyceta | Mucoromycota | Mortierellomycotina | Mortierellomycetes |
| CEJ03135 | PX??? | acyltr. | Rhizopus microsporus | Zygomyceta | Mucoromycota | Mucoromycotina | Mucorales |
| KAG2190366 | PX | hypo | Mucor plumbeus | Zygomyceta | Mucoromycota | Mucoromycotina | Mucorales |
| KAG2189189 | PX | hypo | Umbelopsis vinacea | Zygomyceta | Mucoromycota | Mucoromycotina | Umbelopsidomycetes |
| EWM21801 | PX | PX | Nannochloropsis gaditana | non-fungi | Ochrophyta | | Eustigmatophyceae |
| X6M1E7_RETFI | ? | E2o | Reticulomyxa filosa | non-fungi | Retaria | Foraminifera | incertae sedis |
| X6NR14_RETFI | ? | E2b | Reticulomyxa filosa | non-fungi | Retaria | Foraminifera | incertae sedis |
| OLY84536 | PX | PX | Smittum mucronatum | Zygomyceta | Zoopagomycota | Kickxellomycotina | Harpellales |
| PIA15600 | PX | hypo | Coemansia reveresa | Zygomyceta | Zoopagomycota | Kickxellomycotina | Kickxellomycetes |
| KXN72557 | PX | hypo | Conidiobolus coronatus | Zygomyceta | Zoopagomycota | Entomophthoromycotina | Entomophthoromycetes |

### Supplementary figures

#### Supp. Fig S1

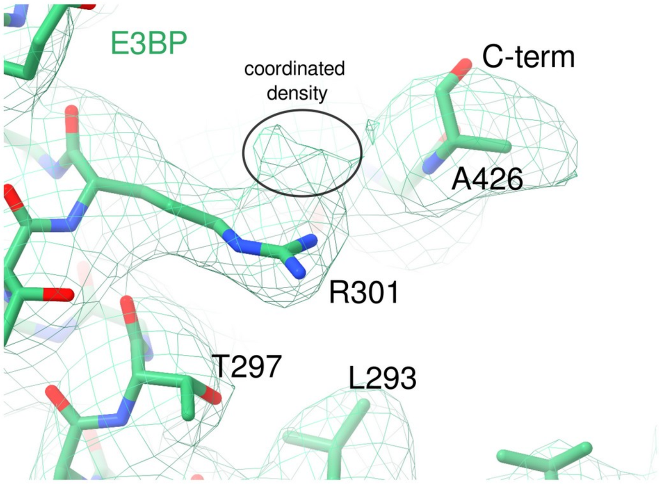

**C-terminal coordination by R301**. R301 i s one of few near-universally conserved residues in fungal E3BP, and seems fully coordinated in the present reconstruction. Given the surrounding hydrophobicity that stabilizes the both the E3BP monomer fold and the homomeric trimer point of contact, the only option for such coordination apears to be the C-terminal carboxyl group of A426. R301 does show additional density to imply such a coordination, however the resolution does not permit unambiguous placement of the 2 C-terminal residues. The penultimate residue of *N.crassa* E3BP is unusual by R425 (not shown), which is almost exclusively small and hydrophobic. This may incur increased flexibility to cause the reduced resolution in the present reconstruction.

#### Supp. Fig S2

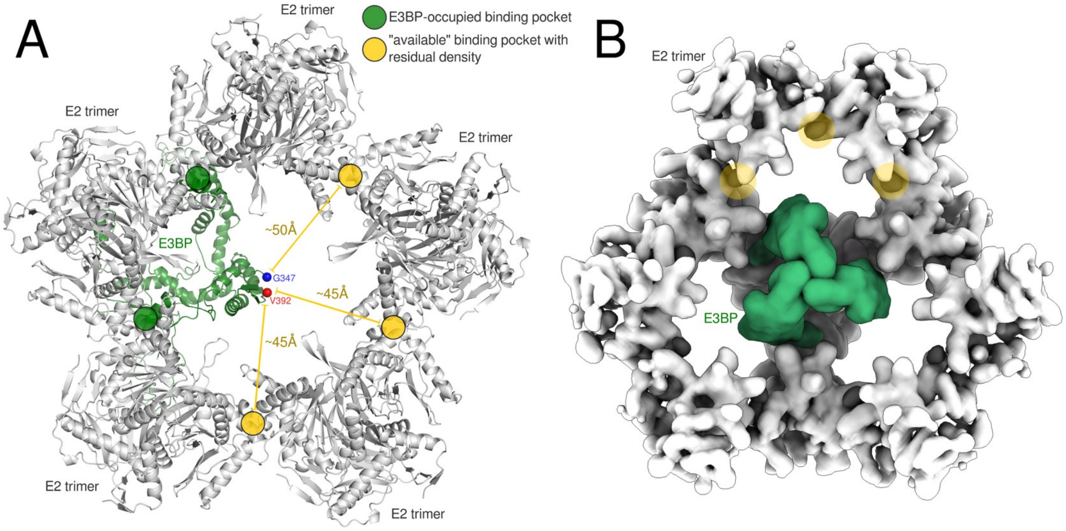

**Available binding sites in E3BP-occupied PDC cores. A** E3BP residues wihtin the range G347-V392 are not resolved in the present reconstruction due to disorder, in line with prediction. The residues however colocalize, approximately to the 5-fold symmetry axis ot he PDC core assembly. The CBD fold here sterically prevents addition E3BP trimers from utilizing ny of the remaining 3 binding sites that are most proximal to both G347 and V392. **B** Residual density is however observed in these interfaces, which is attributed to eithar a) the M3 motif residing in the disordered region between G347 and V392, or b) additional non-trimeric E3BP bound through the M2 motif.

#### Supp. Fig S3

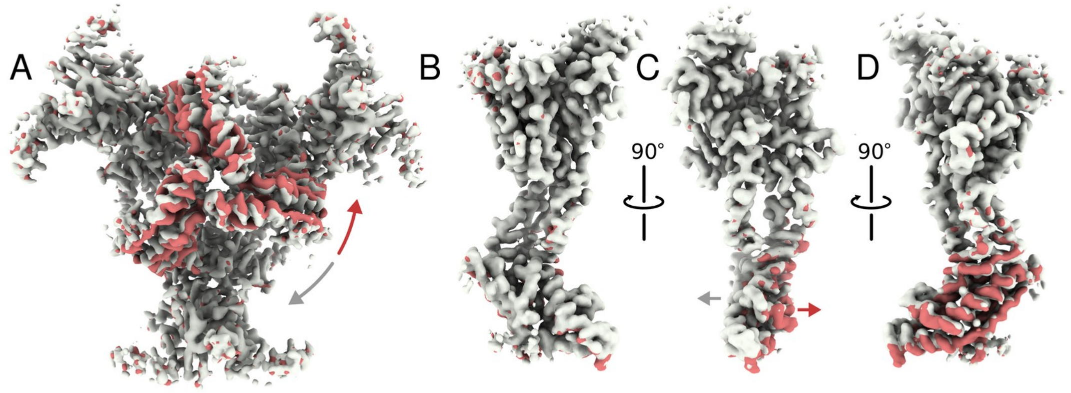

**Flexibility in E3BP trimers. A** Particles of E2 trimers with high occupancy of E3BP were classified into 2 classes (gray and red, respectively) without additional alignments and C3 symmetry. The forced symmetry restricts the classifiable range of motion to rotation around the C3-axis and translation along it, but a view from within the core nevertheless shows a range of flexibility of the CBD with respect to the E2 core that will limit attainable resolution. **B-D** Isolating a single E2 monomer and E3BP monomer shows this  more clearly. Parametrization of this range of motion is expected to improve the resolution further, but this was not conducted in the present work.

### Supp. Fig S4

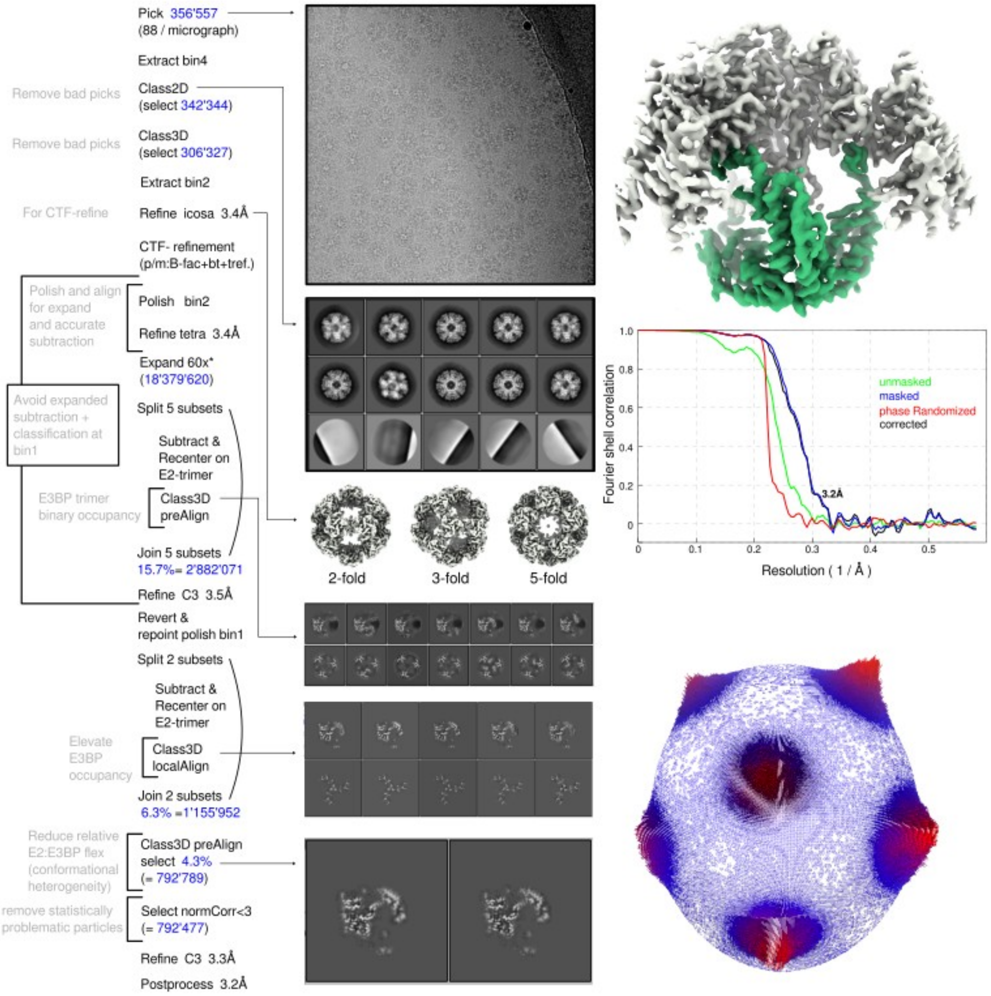

**Cryo-EM data processing details**. The processing pipeline is represented from top to bottom. Particle numbers are highlighted in blue, and percentages are relative to the number following 60x symmetry-expansion. Annotations in gray clarify the motivation of each step. Visual representations of 3D-classifications are by central slice in orthogonal views. The final reconstruction is shown as a surface representation, showing E3BP in green and E2 core in white. Gold-standard FSC is shown, as established through RELIONs post-processing procedure. The orientational distribution following the final 3D-refinement under C3-symmetry shows 4 clusters in the asymmetric unit of the orientational plot, indicating that 5-fold faces are most prevalent in the original data, as is also perceived from micrograph images and 2D-classes. Expansion utilized a custom icosahedral expansion matrix compatible with sym=T to allow easy C3-refinement following re-boxing. relion_symmetry_expand was provided a symmetry file with the relion I2 rotation matrix rotated to match the relion T-symmetry though it's –sym flag:

```
# IcosaCompatWithDefaultTetra.sym
rot_axis    2    0               0.816496        0.57735
rot_axis    3    0               0               1
rot_axis    5    -0.3717480344   0.9091823044    0.1875924741
```

Such a file can in theory be provided to relion_refine, but this an unsupported feature that will not work without further modifications to the code, since the removal of the redundant portion of the orientational sampling space is only defined for the predefined symmetries in RELION. A work-around that will by-pass this issue without fixing it, is to assign C1-symmetry for the sampling but the specified symmetry during reconstruction. This is not recommended. Hence, the present work only utilized predefined symmetries during refinement, and instead defined an expansion symmetry that was compatible with the tetrahedral symmetry.

NOTE: The "revert and re-point polish bin1"-step involves 1) reverting the classified subset of the subtracted and re-centered particles to point to the polished bin2 full-core particles, 2) re-running the previous polishing step and output at bin1, 3) edit the reverted file and relevant fields of the optics group to reference the new polish bin1 job, 4) run a refinement using the edited file starting local searches, and 5) subtracting the polished bin1 particles using the new refinement. In essence, this permits classification at bin2 and subsequent up-scaling to bin1. This was necessary due to the high particle number after expansion in combination with particle subtraction, which generates a new images stack with one particle per alignment, as opposed to one particle per original un-expanded particle.

#### Supp. Fig S5

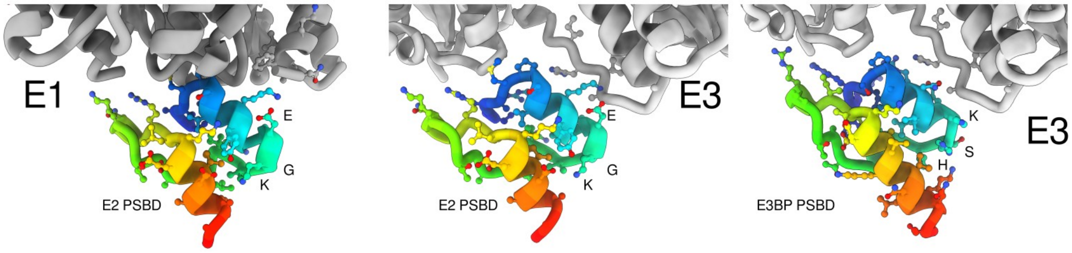

**PSBD binding to peripheral E1 and E3. A** Computational modeling of the human E1-E2 interface. The EKG-motif is indicated, facing away from the E1 binding interface. This motif is conserved in both animal and fungal E2, but lacking in both animal and fungal E3BP (cf. Fig. 4A-B). **B** Superposing the EKG-carrying E2 PSBD onto the E3 binding site reveals that it has possible interactions with an E3 loop. **C** A computational model of the E3-E3BP interface also shows that the EKG motifs if not nearly preserved.

#### Supp. Fig S6

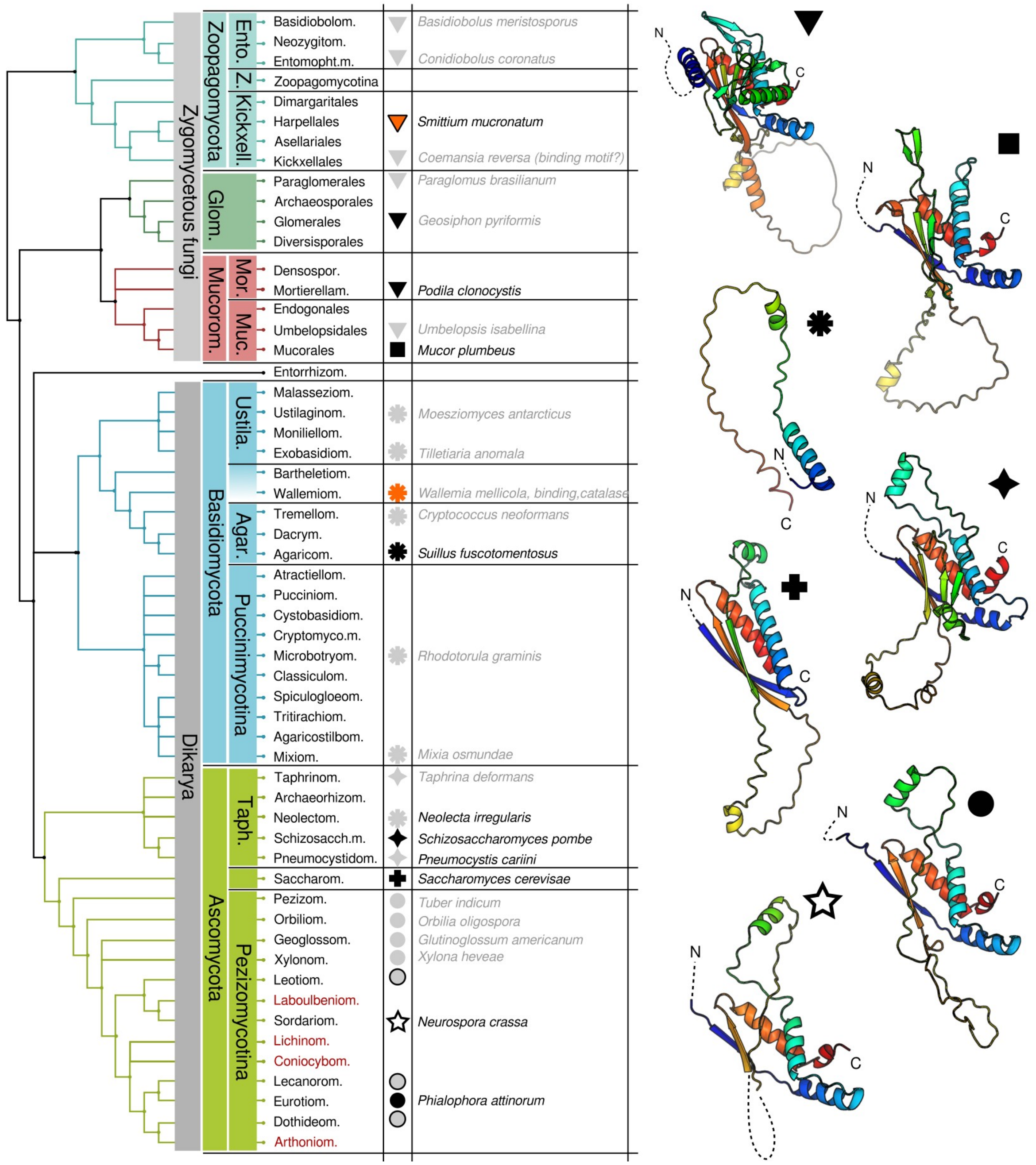

**Phylogenetic relationship of fungal species and E3BP.** The relationship of fungal species is taken from Naranjo-Ortiz & Gabaldón, Biol. Rev. (2019), 94, 2101–2137. Symbols within the table are legends which refer to the most similar E3BP type, as inferred from sequence similarity to models constructed by alphafold2. Black symbols indicate the model shown on the right. A star indicates *N.crassa*, and the structure determined in the present work.

#### Supp. Fig S7

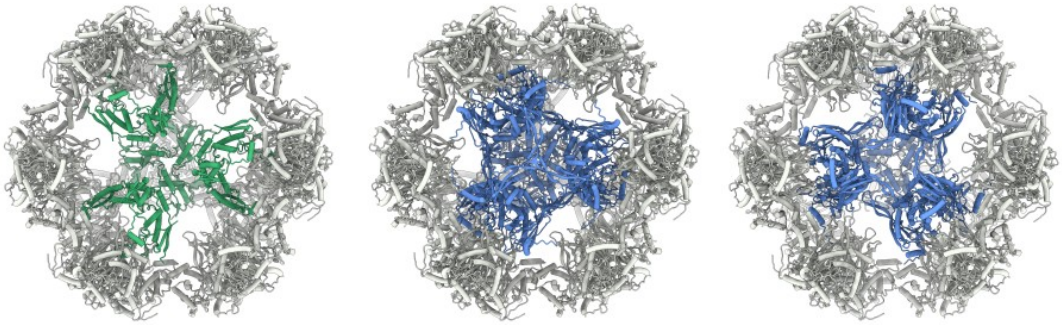

**Comparison of observed and potential core-internal E3BP domains. A** The E3BP CBD permits 12 internal monomers as 4 trimers. **B** Superposing the full ancestral E2 fold onto the observed CBD results in minimal steric clashes, limited to the core-internal loop of the superposed E2 copies. The specific arrangement of the trimers interior to the core however matters. In native preparations of *N.crassa* PDC, it's CBD was shown to arrange in both of two possible core-internal configurations that maximized occupancy to 12 monomers per core. **C** The alternate configuration of core-internal CBD results in further reduced clashes, again limited to the very tip of the core-internal loop of the superposed E2 copies. In this configuration, these loops extend not towards each other, but towards the CBD core fold of each other, producing a degree of potential domain-swapping. It is emphasized that this figure depicts theoretical models under and extended transacetylase fold if it were to occupy a position which aligns with that of the E3BP CBD.
